# Preservation of vision after chemically induced retinal ganglion-like-cell transplantation

**DOI:** 10.1101/2025.11.06.686630

**Authors:** Rana Muhammad Shoaib, Pratigya Tripathi, Sasha Medvidovik, Ryan Lin, Aregnazan Sandrosyan, Biraj Mahato

**Author notes:** To whom correspondence should be addressed: Biraj Mahato, Department of Surgery, Division of Ophthalmology, Children’s Hospital Los Angeles, 4650 Sunset Blvd, Los Angeles, California, CA 90027. These authors contributed equally.

## Abstract

Optic neuropathies are a leading cause of irreversible blindness in children and adults, primarily due to the loss of retinal ganglion cells (RGCs). Currently, there are no effective treatments available to preserve vision in affected patients. Although RGCs derived from pluripotent stem cells present a promising therapeutic strategy, the efficient generation of human RGCs remains a significant challenge. Here, we report a facile method using a combination of five small molecules to reprogram human primary fibroblasts into **C**hemically **I**nduced **R**GC-like **C**ells (CiRGCs), which we term CiRGCs, within just four days. scRNA-Seq analysis revealed that these *in vitro*–generated CiRGCs express canonical RGC markers such as *Pou4f1, Brn3b, Rbpms,* and *Sncg*, as well as several RGC subtype-specific genes. Notably, CiRGCs cluster near native RGCs derived from day 59 fetal retina from published datasets. Furthermore, scATAC-Seq showed open chromatin at RGC-specific promoters and closed chromatin at fibroblast-specific promoters in CiRGCs, confirming successful lineage conversion. Functionally, transplantation of CiRGCs into models of excitotoxic RGC injury led to improved electrophysiological responses for up to 2.5 months possibly through a neuroprotective mechanism. Mechanistically, scRNA-Seq analysis indicated that CiRGC reprogramming proceeds via activation of potential youthful cellular pathways in intermediate cell clusters. In summary, our results establish a rapid, chemical-based strategy for CiRGC reprogramming and highlight its potential as a novel cell therapy approach for treating optic neuropathies, including glaucoma, where RGC loss is the final common pathway.

## Introduction

Optic neuropathies encompass a group of disorders characterized by dysfunction and degeneration of the optic nerve, ultimately leading to optic atrophy and irreversible vision loss. A hallmark of these conditions is the progressive loss of RGCs and their axons, a common feature in both acquired diseases such as glaucoma, inherited disorders like Leber’s Hereditary Optic Neuropathy (LHON) and optic nerve hypoplasia ^1–3^. In glaucoma, elevated intraocular pressure (IOP) and aging are major risk factors contributing to RGC degeneration, while LHON is primarily driven by mitochondrial DNA mutations affecting RGC survival^4^. Despite significant global prevalence across both adult and pediatric populations, effective therapies that restore vision in these patients remain unavailable. Developing methods to generate functional RGCs holds significant potential for advancing regenerative medicine, disease modeling, and high-throughput drug discovery for optic neuropathies. While protocols employing embryonic stem (ES) cells and induced pluripotent stem (iPS) cells have demonstrated the ability to differentiate into RGC-like cells, these approaches are technically complex, time-consuming, and often yield heterogeneous populations—posing significant barriers to clinical translation^5^. Recently, direct reprogramming using small molecules has recently emerged as a powerful and more accessible strategy for generating functional cell types, including neurons, astrocytes, cardiomyocytes, photoreceptors, and even pluripotent stem cells^6–11^. However, direct pharmacological reprogramming to RGCs has not been realized. Here we have established a small molecule-based protocol capable of inducing human fibroblasts into CiRGCs in just 4 days. Transcriptome and chromatin analysis indicated that these cells are neurons containing RGC-like gene expression and chromatin signature. 5C-induced chemical reprogramming approach may serve as a novel therapeutic strategy for glaucoma and other optic neuropathies where RGC death is the final common endpoint.

## Results

### 5C reprograms human primary fibroblasts into CiRGCs

A prior study demonstrated a combination of five small molecules (valproic acid, CHIR99021, RepSox, Forskolin, and IWR1) that successfully reprogrammed fibroblasts into photoreceptors, leading to restored vision upon subretinal transplantation in blind mice^12^. Observing PAX6^+^ progenitor-like cells at an earlier stage during this reprogramming process^13^, we hypothesized that modifying this cocktail might enable the generation of RGC like neurons. Subsequently we modified the cocktail by replacing IWR1 (that induces photoreceptor lineage) with ISX9 considering its role in inducing *POU4F1* expression, a marker gene for RGCs^14^. We generated human lung fibroblast (HLF) derived CiRGCs using this new combination in just 4 days (**Figure 1A**). CiRGCs from lung fibroblasts also express RGC marker proteins POU4F1 and SNCG (**Figure 1B**). To further characterize CiRGCs, we performed scRNA-Seq for chemically reprogrammed cell population. A total of approx. 8000 cells were included in bioinformatic analysis that were divided into 8 distinct clusters with clusters 6 and 7 separated from rest of the clusters (**Figure 1C**). Cluster-wise analysis for *Pou4f1⁺Rbpms⁺* cells—the two most reliable markers of RGCs^32^—showed that CiRGCs were predominantly distributed across all clusters except clusters 5, 6, and 7 (**Figure 1D**). A similar distribution pattern was observed for *Sncg⁺Nefm⁺Nefl⁺* CiRGCs (**Figure 1E**). Heatmap analysis further revealed two clusters containing both *Pou4f1⁺* and *Pou4f2⁺* CiRGC subtypes (**Figure 1F**). Specifically, cluster 5 expressed RGC-defining genes including *Pou4f2, Sncg, Nefm, Nefl, Isl1, Gap43, Tubb3,* and *Thy1*, whereas cluster 6 expressed *Pou4f1, Rbpms, Isl2, Elavl3, Elavl4,* and *Sox4*. Notably, neither of these clusters expressed fibroblast-associated genes *Col1a2* and *Axl*, suggesting complete conversion to the retinal neuronal lineage. In contrast, cluster 4 cells expressed the fibroblast marker *Col1a1* along with low levels of RGC-specific genes, indicating a potential intermediate state during reprogramming.

**Figure 1:**
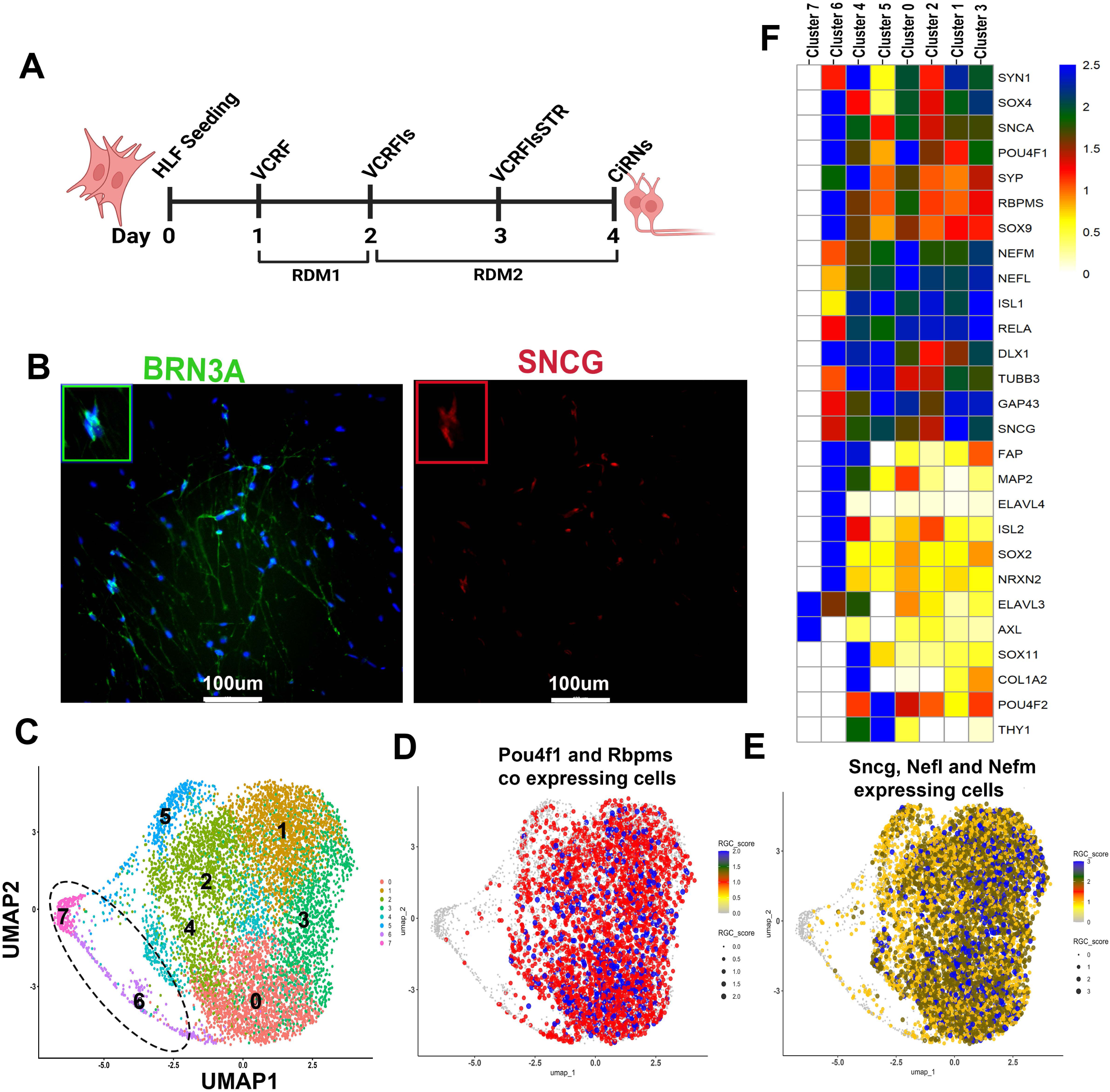
Reprogramming human lung fibroblasts (HLF) to Fd-CiRGCs. (**A**) Scheme for the reprogramming experiment. V; Valproic acid, C; CHIR99021, R: Repsox; F: Forskolin; Is: ISX9; S: Shh; T: Taurine, R: Retinoic acid; RDM; RGC differentiation medium. (**B**) Immunofluorescence staining of CiRGCs on day 4, expressing POU4F1 and SNCG. (**C**) UMAP analysis indicated 8 distinct clusters, and two separate clusters (possible intermediate cells) indicated in circle. (**D**) Expression of RGC specific genes in various clusters. (**E-H**) Expression of indicated RGC specific genes in POU4F1, POU4F2^+^, POU4F1^+^RBPMS^+^, SNCG^+^NEFM^+^NEFL^+^ cell sets.

Furthermore, we generated specific cell subsets from the total population based on the expression of *Pou4f1⁺*, *Sncg⁺*, and *Sncg⁺Nefl⁺Nefm⁺* gene combinations. Analysis of these subsets revealed that the same cells co-express additional canonical RGC-specifying genes. The *Pou4f1⁺* subset expressed multiple RGC markers, including *Gap43, Isl1, Isl2, Rbpms, Sncg, Sox4, Nefl,* and *Tubb3* (**Figure S1A, B**). Similarly, the *Sncg⁺* subset showed co-expression of *Pou4f1, Gap43, Sox4, Nefl, Isl1, Isl2,* and *Rbpms* (**Figure S1C, D)**. Consistent with this pattern, the *Sncg⁺Nefl⁺Nefm⁺* subset co-expressed *Pou4f1, Isl1, Isl2, Rbpms, Gap43, Sox4,* and *Tubb3* (**Figure S2A-C)**. Collectively, these findings indicate that the 5C treatment can reprogram human lung fibroblasts into RGC-like neurons.

### CiRGCs show similarity to day 59 human fetal RGCs

We compared single-cell gene expression signatures between CiRGCs and human fetal retina day 59 utilizing a previously published dataset^15^. UMAP analysis was performed between CiRGCs from lung fibroblasts and fetal retina at day 59 after integration of both data sets concurrently (**Figure 2A**). We found a total of 14 clusters in the UMAP analysis when both datasets are embedded and analyzed concurrently. Cluster analysis indicated clusters 9, 4 and 13 corresponding to the CiRGC data set fall near day 59 fetal retinal cells, rest are separated (**Figure 2B**). Subsequently unbiased cluster annotation reveals that CiRGC population containing various cell types (including progenitors, photoreceptors etc.) and an unknown population corresponding to clusters 9 and 4 which was situated closely to the RGC cluster (#2, native RGCs) from the d59 fetal retina (**Figure 2C**). We hypothesized cells in these unknown clusters are RGC like cells and subsequently we compared these clusters with native RGCs from fetal retina present in cluster 2. Heatmap analysis indicated expressions of all major RGC genes such as *Pou4f1, Pou4f2, Sox4, Isl1, Sncg, Nefl, Nefm, Tubb3, Gap43* clusters 9, 4 when compared to native RGCs in cluster 2 (**Figure 2D**). Next, we performed unbiased marker gene analysis by commanding the program to find out the most abundant 20 genes in each cluster. Analysis revealed many similar regions in the heatmap that include RGC specific genes such as *Gap43, Sncg, Isl1* etc (**Figure 2E green boxes**). These results suggest that chemically induced unknown cell population is RGC like neurons and fall near native RGCs.

**Figure 2:**
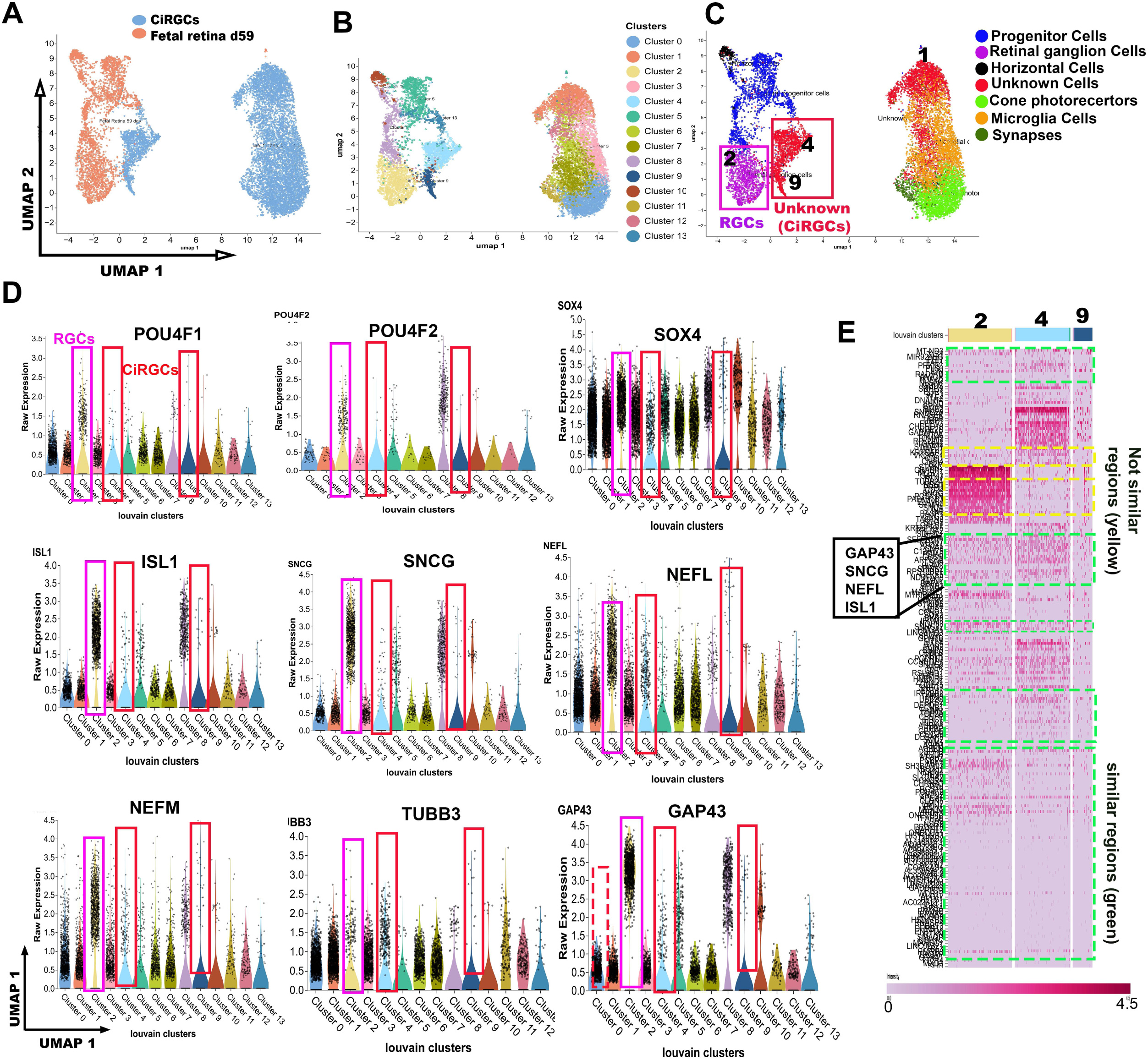
Comparison between fibroblast derived CiRGCs and native RGCs from fetal retina d59. **(A)** UMAP analysis after CiRGC and fetal retina d59 after embedding two datasets. (**B**) Total of 14 clusters were found in UMAP after embedding. (**C**) Unbiased cell type annotation showing retinal progenitors, cone photoreceptors, synapse cells and an unknown cell in CiRGC population. Unknown cells located in clusters 9 and 4, fall near native RGCs from day 59 fetal retina in cluster 2. (**D**) Cluster wise comparison of indicated RGC genes expression. Purple: native RGCs; Red: CiRGCs; Dashed Red: possible immature intermediates. (**E**) Unbiased marker gene analysis (20 genes per cluster) of the indicated clusters by heatmap showing presence of key RGC genes. Green box: similar regions. Yellow: differential region.

### CiRGCs exhibit similar diversity when compared to day 59 fetal RGCs

RGCs exhibit remarkable diversity, comprising various subtypes^16^. We investigated subtype-specific gene expression in CiRGCs and in human fetal retina on day 59 using *Pou4f1*^+^ and *Sncg*^+^*Nefm*^+^*Nefl*^+^ cell sets. These subtype specific genes were selected from published studies and covers all 40 subtypes^16–18^. Our analysis revealed that majority of the subtype specific gene are expressed in similar levels between CiRGCs and fetal retina day 59 (**Figure S3A and B**). For example, transcription factors *Runx1* and *Fst* uniquely enriched in subtype 27 are present in both CiRGCs but not in fetal retina day 59 (**Figure S3A and B**; green arrows). *Trhr* (subtype 31), *Pde1a* (subtype 7) and *Bcl11b* (subtype 11) also present in CiRGCs (**Figure S3A and B;** red arrows). Additionally, RGCs are divided into midget cells, which have smaller soma size, and the larger size parasol cells. We have analyzed these subtype specific gene expression in *Pou4f1*^+^ cell sets between native RGCs (cluster 2) and CiRGCs (cluster 4) from figure 2C. The results revealed that CiRGCs express OFF parasol RGC markers, while some of these markers were absent in fetal cells (**Figure S4A and B**). Interestingly, approximately 5% of the *Pou4f1+* CiRGCs show expression of direction selective RGC marker *Fstl4,* indicating reprogramming preference to this subtype. We also noted absence of *Zic1* expressing subtype 34 from CiRGC population (**Figure S4C;** red and green arrows). RGCs. These results suggest CiRGCs express RGC subtype genes which to RGCs from day 59 fetal retina.

### CiRGCs contain open chromatins in RGC specific promoters

Next, we analyzed the global chromatin landscape of CiRGCs using single-cell ATAC sequencing (scATAC-Seq). A total of approx. 20,000 nuclei were included in the analysis. UMAP clustering based on the *promoter sum* feature revealed seven major clusters (**Figure 3A**). The promoter sum for a given gene was determined by the number of transposases cut sites (peaks) per cell within the promoter region (−1 kb to +100 bp from the transcription start site). A higher promoter sum reflects greater promoter accessibility and more open chromatin. Notably, Cluster 4 was separated from the remaining clusters, leading us to hypothesize that this cluster represents CiRGCs. Cluster-wise heatmap analysis revealed that Cluster 4 exhibited open chromatin regions in all major RGC-specific promoters, including *POU4F1, SNCG, NEFM, NEFL, ISL1, ELAVL3, GAP43, POU4F2, SOX4,* and *ISL2*. In contrast, the fibroblast gene *COL1A2* displayed a closed promoter configuration, indicating complete cellular conversion (**Figure 3B**). A global heatmap analysis across all nuclei further confirmed open promoter accessibility in canonical RGC-specifying gene promoters (**Figure 3C**). Moreover, cell set heatmap analysis restricted to *POU4F1+* cells—similar to our scRNA-Seq approach—showed that cells with open *POU4F1* promoters also exhibited open chromatin at other canonical RGC gene promoters (**Figure 3D**).

**Figure 3:**
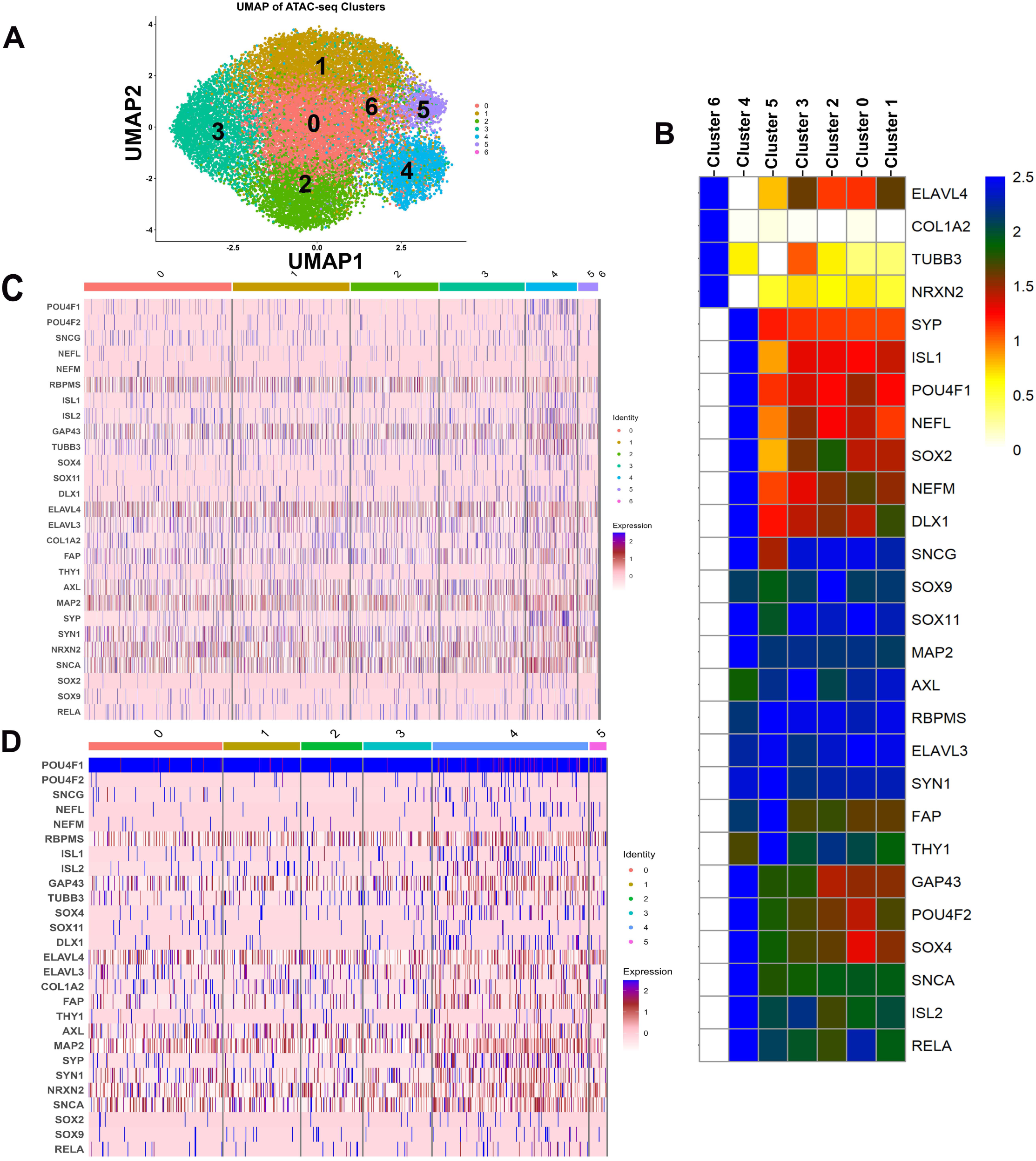
scATAC-Seq analysis of CiRGCs indicated open chromatin regions in the promoters of major RGC genes and the results are consistent with the scRNA-Seq data. **(A)** UMAP analysis of approx. 20000 cells showing 7 clusters. (**B**) Heatmap analysis showing cluster wise open chromatins in the promoter regions of the indicated RGC specifying genes. (**C**) Heatmap showing clusterwise opening of chromatins in the indicated genes. (**D**) Heatmap showing POU4F1^+^ cell set expressing canonical RGC specific genes.

Subsequently, we aligned the scATAC-Seq reads to annotate genomic feature distributions, which included enrichment across transcription start sites (TSS), transcription termination sites (TTS), introns, intergenic regions, and untranslated regions (UTRs) (**Figure 4A**). We then focused on promoter accessibility in RGC-specifying genes that were upregulated in our scRNA-Seq dataset. Nuclei were filtered by *POU4F1* promoter sum using the algorithm’s barcode filtering function. Open chromatin regions were visualized in *POU4F1* promoter–positive cells using the peak viewer tool, which displays peak size and the proportion of cells containing those peaks. Consistent with our scRNA-Seq results, prominent peaks indicating accessible chromatin were observed in RGC gene promoters such as *POU4F1, ISL1, POU4F2, SNCG, NEFL,* and *NEFM* (**Figure 4B–G**). In contrast, fibroblast gene promoter *COL1A2* lacked significant accessibility, confirming loss of fibroblast identity (**Figure 4H**).

**Figure 4:**
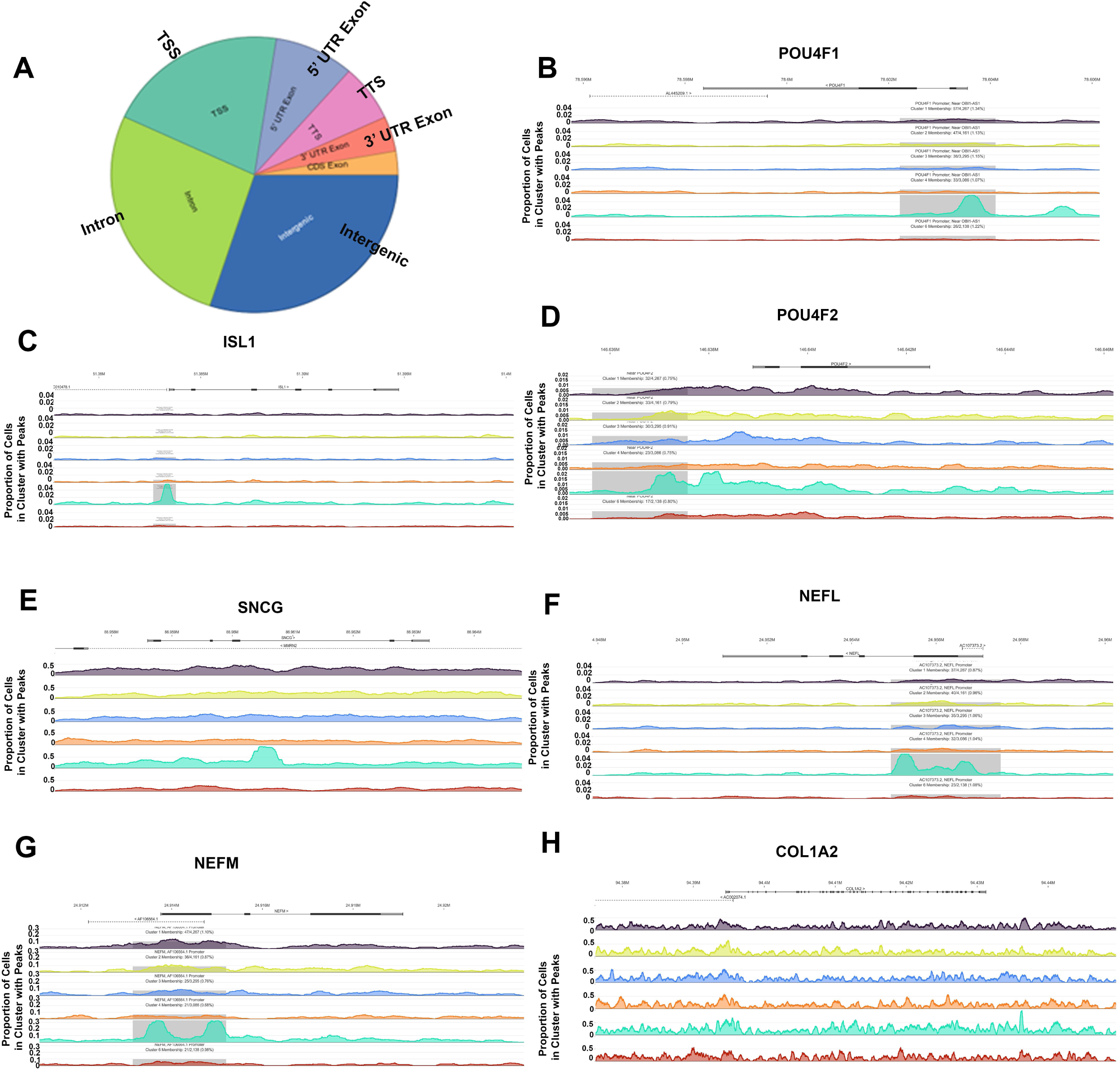
scATAC-Seq analysis showing gene section breakdown and open chromatin peaks in the promoter regions of the indicated genes. (**A**) Gene section breakdown indicated presence of various regulatory regions in the sequenced data. (**B-H**) Promoter peak analysis indicated open chromatins in the indicated gene promoters.

Together, these findings demonstrate that CiRGCs possess open chromatin structures at RGC-specific promoters while maintaining closed configurations at fibroblast-specific promoters, indicative of successful and complete cellular reprogramming.

### CiRGC reprogramming proceeds through a younger intermediate cellular stage

To investigate the mechanism underlying CiRGC reprogramming, we hypothesized— chemical reprogramming can revert somatic cells into a younger partially reprogrammed state without inducing full pluripotency based on a prior study^20^. Cluster wise pathway enrichment analysis revealed significant upregulation of genes involved in oxidative phosphorylation, mitochondrial electron transport, and mTOR signaling, along with downregulation of inflammatory pathways, including interferon alpha/gamma responses, complement cascade, and JAK/STAT signaling similar to a younger progenitor stage (**Figure 5A**). These transcriptional changes closely parallel those observed in previous studies where small molecule cocktails were shown to reverse cellular aging phenotypes^19^. These findings support a model in which CiRGC reprogramming proceeds through a metabolically active, progenitor-like intermediate state characterized by reduced inflammatory signaling and enhanced mitochondrial function (**Figure 5B**).

**Figure 5:**
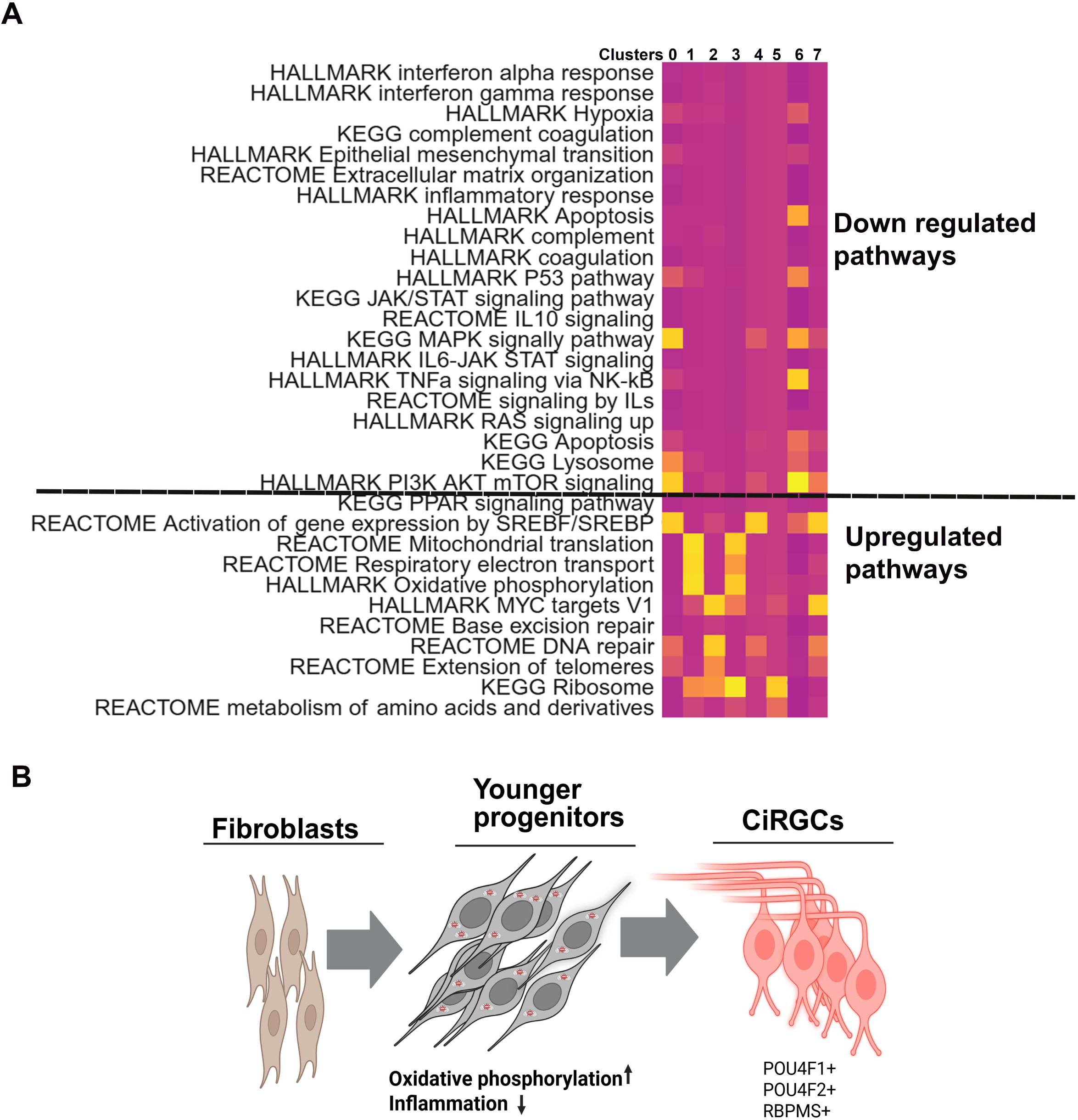
Enhanced younger cellular pathways during CiRGC reprogramming. (A, (A) Enrichment of indicated youthful cellular pathways involved in the intermediate clusters during CiRGC reprogramming. (**B**) Diagram showing CiRGC reprogramming through an intermediate younger cellular stage.

### Transplanted CiRGCs improve visual functions after NMDA induced injury

Next, we assessed the therapeutic potential of CiRGCs by intravitreally transplanting them into NMDA-mediated RGC injured mice. First, we generated a BRN3B-promoter reporter using a promoter-less GFP vector (Clontech), based on a previous study (**Figure S5**)^20^. Human fibroblasts were transfected with BRN3B-GFP lentivirus and reprogrammed using the 5C protocol described above (**Figure S6)**. For functional studies, we injected NMDA (100 mM, 1.5 µL/eye) into 8–10-week-old C57BL/6 mice (**Figure 6A**). Seven days after NMDA treatment, we assessed retinal and visual function using pattern ERG (pERG) and pattern VEP (pVEP). Mice with diminished pERG/pVEP responses were selected for transplantation. We transplanted 200,000 CiRGCs per eye, following MACS separation using fibroblast-specific magnetic beads and FACS separation. pERG and pVEP were recorded at the time points indicated in Figure 6A. We observed a gradual increase in pERG amplitude starting from day 7 and continuing through day 30 (**Figure 6B**). From day 60 onward, pERG amplitude began to decline, although it remained significantly higher than the post-NMDA baseline (**Figure 6B, right bar graphs**). In contrast, pVEP amplitude gradually increased up to day 60 and then showed a moderate decline by day 75 but remained significantly improved compared to post-NMDA levels. Control mice injected with PBS showed no improvement in either pERG or pVEP (**Figure 6C**).

**Figure 6:**
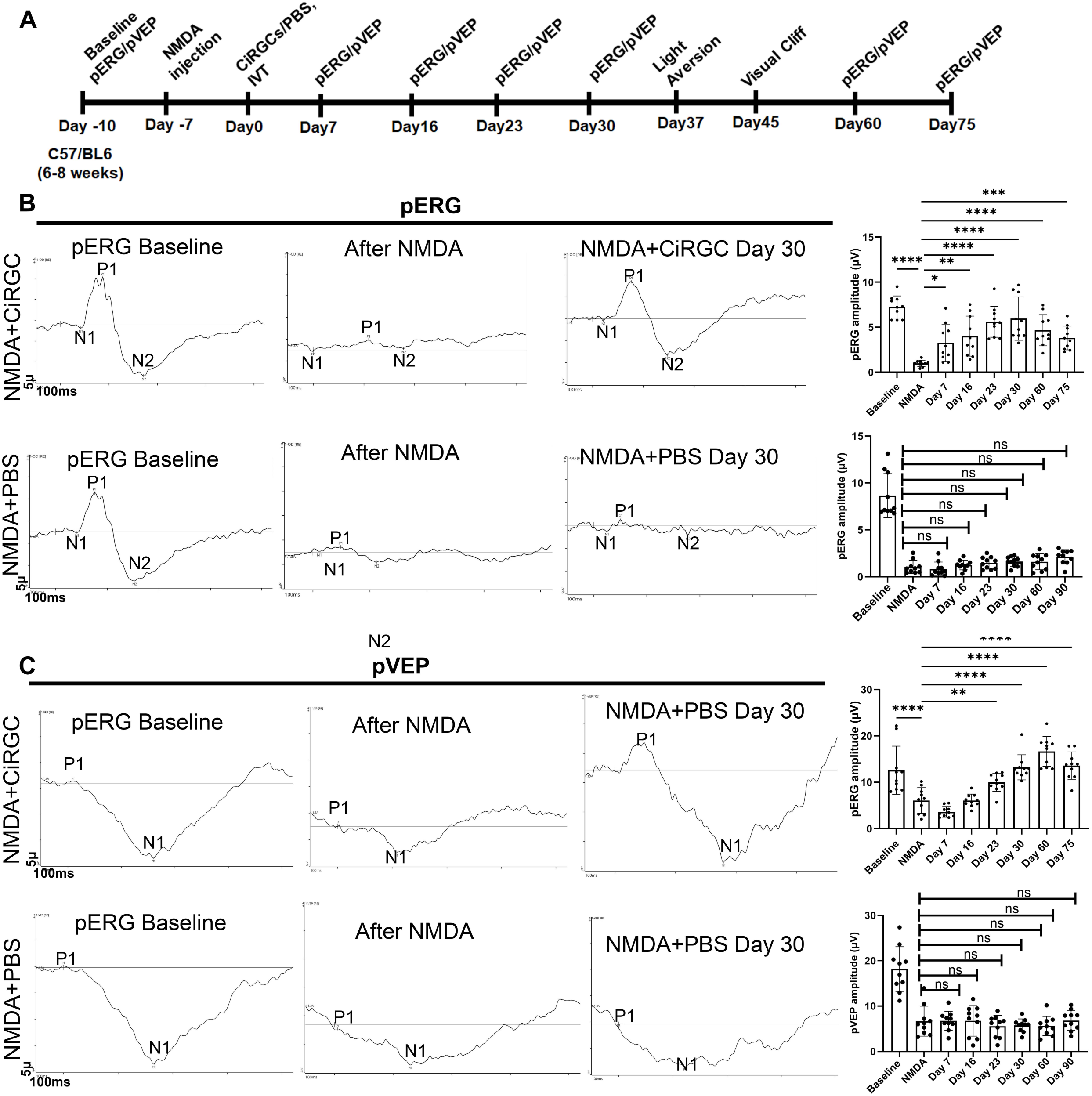
Improvement of electrophysiological functions (pERG/pVEP) after CiRGC transplantation in NMDA injures mice. **(A)** Timeline for the transplantation experiment. (B) Right: Representative waveforms for pERG from CiRGC and PBS injected eyes. Right: pERG amplitudes are plotted for CiRGC and PBS eyes. (n=5 animals for each group tested). (**C**) Right: Representative waveforms for pVEP from CiRGC and PBS injected eyes. Right: pVEP amplitudes are plotted for CiRGC and PBS eyes. (n=5 animals for each group tested).

We next conducted a longitudinal light avoidance test in NMDA-injured mice that exhibited improved pERG/pVEP responses, to assess whether enhanced retinal function correlates with the restoration of visually evoked behaviour^21–23^. This test relies on the innate tendency of mice to avoid brightly lit areas. Animals were placed in a light/dark box for 300 seconds, and the time spent in the brightly illuminated compartment (2800 lux, approximating a cloudy outdoor environment) was recorded. Wild-type mice preferred the dark side (194 ± 14 sec), whereas PBS-treated NMDA-injured mice showed a reduced aversion to light (123 ± 12 sec). In contrast, CiRGC-transplanted mice demonstrated a significant preference for the dark compartment (186 ± 23 sec), indicating improved light avoidance behavior (**Figure 7A and B**). Similarly, the latency to first entry into the dark chamber was comparable between wild-type (8 ± 1.5 sec) and CiRGC-treated mice (7 ± 3.9 sec), while PBS-treated NMDA-injured mice exhibited a significantly delayed response (38 ± 5 sec) (**Figure 7C**). We next performed a longitudinal visual cliff test, a behavioral method used to evaluate depth perception and the fear of crossing the deep side of a platform^24–26^. Wild-type mice typically show a strong innate tendency to avoid the deep side of a visual cliff, favoring the shallow side. In this test, mice were placed on a central platform between the deep and shallow sides, and their choices were carefully recorded. As expected, wild-type mice demonstrated a clear preference for the shallow side, with 96 ± 3.9% of choices. In contrast, PBS-treated NMDA-injured mice showed a marked impairment in depth perception, with only 32 ± 8.9% of choices to the shallow side. Conversely, the CiRGC treated group demonstrated a more favorable outcome, with 56±8.94% of choices directed towards the shallow side in NMDA injured mice (**Figure 7D and E**). In assessing the cumulative duration of stay on both the shallow and deep sides, it became evident that wild-type mice exhibited a distinct preference towards the shallow side. Conversely, the PBS treated NMDA injured mice showed no clear preference, indicating a compromised performance. Notably, CiRGC-transplanted mice showed a preference for the shallow side (**Figure 7F**). These findings strongly indicate a recovery of long-term visual functions in RGC injured mice after CiRGC transplantation. At this stage (day 75), we hypothesized that the gradual decline in pERG amplitude could be due to immune-mediated rejection. To investigate this, we sacrificed the mice and examined the retina for the presence of transplanted cells. We detected HuNu⁺/RBPMS⁺ and GFP⁺/RBPMS⁺ cells in the transplanted retinas (**Figure S7A-D**). The quantification showed the presence of avg 36± 5.8 CiRGCs per eye on day 75 (**Figure S8A**). These results suggest that transplanted CiRGCs can transiently improve visual function after NMDA-induced injury, with therapeutic effects lasting up to approximately 2.5 months.

**Figure 7:**
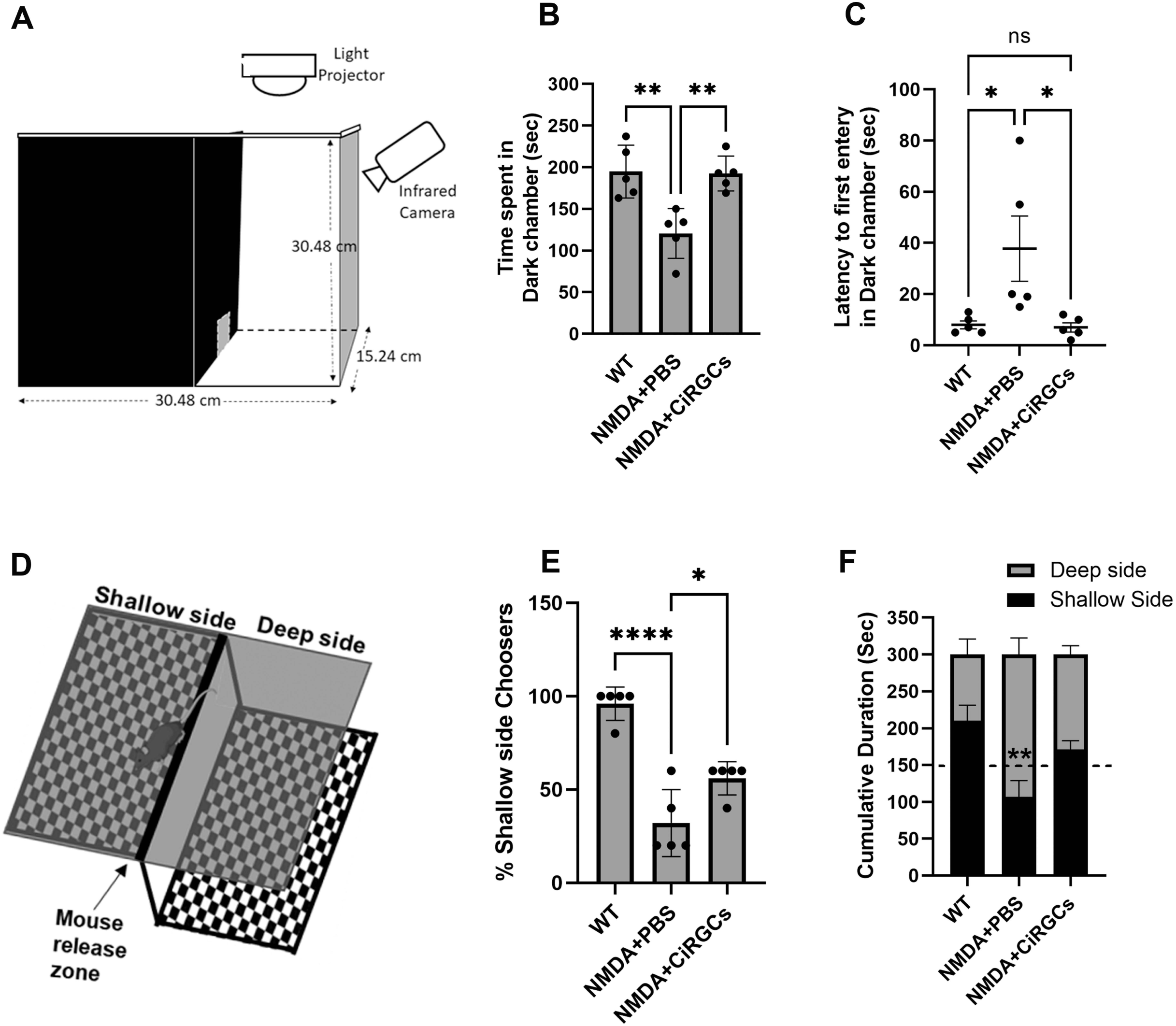
Improvement of light aversion and depth perception behaviors following CiRGC transplantation in NMDA-Injured C57BL/6 Mice. **(A)** Schematic illustration of the experimental setup for the light aversion behavior test. **(B)** Quantification of the average time (± SEM) spent in the dark chamber by each experimental group. (**C**) Quantification of the average latency (± SEM) to first entry into the dark chamber across groups. (**D)** Schematic of the setup for the depth perception (visual cliff) test. **(E)** Percentage of trials in which animals chose the shallow side; five trials were conducted per animal. **(F)** Cumulative duration spent on the deep versus shallow sides after the cliff barrier was removed, shown for each group. Statistical analysis was performed using one-way ANOVA for panels b, c, and e, and two-way ANOVA for panel f.

## Discussions

Here, we report a chemical-only reprogramming strategy that converts human primary lung fibroblasts into RGC-like neurons, offering a potential allogeneic cell therapy approach for glaucoma and other optic neuropathies. This CiRGC reprogramming method was developed by modifying a previously reported chemical cocktail used to generate photoreceptor-like cells^12^. By replacing IWR1 with ISX9 in the cocktail, we successfully obtained CiRGCs; however, a subset of cells retained their photoreceptor identity along with other retinal cell types (**Figure 2C**), suggesting that CiRGC generation may involve shared developmental pathways and a common intermediate progenitor like state. scRNA-Seq analysis identifies three distinct clusters (clusters 9, 4) of unknown cells that express RGC specifying genes (**Figure 2C, D**). Intermediate clusters also showed enrichment for pathways associated with younger cellular stages similar to a previously published study^19^, suggesting that CiRGC reprogramming may proceed through a progenitor-like, younger cellular state (**Figure 5B**). Transcriptome analysis indicated that CiRGCs express vision-forming and pan RGC markers *POU4F1* and *RBPMS*, which are reported to be the best RGC markers across species^27–31^. While expressing essential markers, CiRGCs lacking certain mature RGC-specific markers (such as *THY1*) indicating that these cells are immature and that would be beneficial for cell transplantation studies. Approx. 5% of POU4F1^+^ CiRGCs express direction sensing RGC marker *Fstl4*, a subtype capable of responding to preferred directional motion in response to light or dark stimuli. Intriguingly, CiRGCs exhibited expression of *Isl2* (**Figure 1F**) which is an exclusive marker for contralateral RGCs (cRGCs) that cross their axons in the midline of the optic chiasm and continues to the brain in the contralateral optic tract^32^. Furthermore, CiRGCs express parasol RGC markers which are predominantly responsible for conveying inputs from the eye to the lateral geniculate nucleus in the brain. These molecular characteristics support the potential functional relevance of CiRGCs in vision restoration strategies. Additionally, larger RGCs expressing *Pou4f1* are preferentially susceptible to death in human glaucoma, suggesting *in vitro* derived CiRGCs would be suitable for therapeutic interventions^33^. In our proof-of-concept transplantation study, we utilized the CiRGCs to rescue vision transiently after excitotoxic RGC injury. Since we did not observe efficient CiRGC survival after day30 (however electrophysiology improvement remains significant), we anticipate that CiRGC mediated host RGC neuroprotection may be a possible mechanism for the vision rescue similar to demonstrated published studies in retina and other part of the central nervous system (**Figure S8B**)^15,34,35^. We anticipate CiRGC induced paracrine mechanism of neuroprotection in host RGCs may be attributed to the observed visual function improvement. Further investigation is required to elucidate the underlying processes.

In summary, this study presents a chemical-only method for generating RGC-like neurons from human fibroblasts, highlighting a promising direction for developing cell-based neuroprotective therapies for optic neuropathies associated with RGC loss.

## MATERIALS AND METHODS

### Generation of chemically induced Retinal ganglion cells *in vitro*

Human primary lung fibroblasts (Coriell Institute, Cat# GM05389) were seeded in 24-well plates coated with 0.1% gelatin and cultured overnight. The following day, the normal growth medium was replaced with Retinal Differentiation Medium (RDM1, composition detailed in Suppl. Tables 1), containing small molecules V (0.5 mM), C (4.8 μM), R (2 μM), and F (10 μM). On day 2, ISX9 (Is) (20 μM) along with VCRF (same concentrations as before) were added to the cells in RDM2 medium. On day 3, Taurine (T), Retinoic Acid (R), and Sonic Hedgehog (S) were also added, in conjunction with the other small molecules, in RDM2 medium. On day 4, cells were collected for molecular assays, sequencing studies, and immunofluorescence staining.

### Intravitreal Injection and CiRGC Transplantation

All intravitreal injections were performed under ketamine (85 mg/kg) and xylazine (14 mg/kg, intraperitoneally) anesthesia at the CHLA animal care facility. For the NMDA-induced retinal injury model, 6–8-week-old mice received intravitreal injections of NMDA (Sigma-Aldrich) prepared as a 100 mM solution in PBS. Under a dissection microscope positioned inside a BSL-II laminar flow hood, pupils were dilated with 1% tropicamide prior to injection. Using a 10 μL Hamilton syringe fitted with a 30-gauge needle (Small Hub RN Needle, Hamilton), 1.5 μL of the NMDA solution was injected into the vitreous chamber at a 45° angle relative to the lens.

For transplantation experiments, human fetal lung fibroblasts (passage 6) were transduced with Lenti-Brn3b-GFP (MOI 8) and reprogrammed into chemically induced retinal ganglion cells (CiRGCs). The reprogrammed population was subjected to fluorescence-activated cell sorting (FACS) to isolate GFP⁺ cells, which were then resuspended in RDM2 medium. At 7 days post-NMDA treatment, approximately 2 × 10⁵ GFP⁺ CiRGCs were injected intravitreally under microscopic guidance (Amscope) using a 32-gauge Hamilton microsyringe at a 45° angle to the lens. The needle was held in place for several seconds post-injection to ensure complete release of the cell suspension before being withdrawn slowly. Following injection, the eyelid was repositioned, a drop of antibiotic ointment was applied, and mice were maintained on a 37°C warming pad until full recovery.

### Immunostaining of Tissue Sections and Cultured Cells

Retinal Tissue Sections: Following CO₂-mediated euthanasia, eyes were enucleated and fixed in 4% paraformaldehyde (PFA) prepared in phosphate-buffered saline (PBS) for 15 minutes. After fixation, the cornea was removed, and the resulting eye cups were sequentially cryoprotected by incubation in 15% sucrose, followed by 30% sucrose, overnight at 4°C. The eye cups were then embedded in optimal cutting temperature (O.C.T.) compound and cryosectioned at a thickness of 30 μm.

For immunostaining, cryosections were first washed twice with PBS, followed by three washes with 0.1% PBST (PBS with 0.1% Tween-20). Sections were then permeabilized with 0.25% PBST for 15 minutes. After blocking for 1 hour at room temperature, sections were incubated with primary antibodies overnight at 4°C. The following morning, sections were washed five times with 0.1% PBST and incubated with Alexa Fluor-conjugated secondary antibodies (Alexa 488 or Alexa 594) for 90 minutes. Finally, slides were washed ten times with 0.1% PBST, mounted with Fluoromount-G containing DAPI, and allowed to dry for two hours before imaging.

Cultured Cells (Reprogrammed CiRGCs): For immunofluorescence staining of reprogrammed retinal ganglion cells (RGCs), cultured cells were fixed with 4% PFA in PBS for 20–25 minutes and permeabilized using 0.25% PBST for 30 minutes. Cells were then blocked for 1 hour at room temperature and incubated with primary antibodies (see Extended Data Table 2) overnight at 4°C. The next morning, cells were washed five times with 0.1% PBST and incubated with appropriate secondary antibodies for 90 minutes. After washing with 0.1% PBST, cells were incubated with DAPI solution (1 µg/mL) for nuclear staining. Fluorescent images were acquired using a Leica DMi8 confocal microscope at the CHLA Imaging Core Facility. Quantification of immunostained cells was performed using ImageJ and Adobe Photoshop. For regenerated axon quantification, every fifth tissue section was analyzed, with four sections included per eye (n = 3 eyes).

### RT-PCR and Quantitative real time PCR

RNA samples were extracted by using RNA isolation kit from Zymo Research (MicroPrep R1050). cDNA syntheses were performed by High-Capacity cDNA Reverse Transcription kit from Applied Biosystems according to the manufacturer’s instructions. RT-PCR and qPCR was performed by using specific set of primers listed in thes table 3. Thermal cycler from Applied Biosystems were used for RT-PCR analysis. For quantitative real time PCR, sample were processed using and OneStep Plus real time PCR. Results were normalized to the housekeeping gene Gapdh and analyzed using the ΔΔCt method. Results were presented in fold changes compared to unconverted cells.

### Single cell RNA Sequencing

Single-cell RNA-seq libraries were generated using the 10x Genomics Chromium Single Cell platform and processed with Cell Ranger (v7.1.0) to produce filtered feature-barcode matrices, which were imported into R *(v4.5.1)* and processed using Seurat *(v5.3).* Quality control metrics, including the number of detected genes per cell, total counts per cell, and percentage of mitochondrial reads, were computed to exclude low-quality cells. Normalization, identification of highly variable features, and scaling were performed with NormalizeData, FindVariableFeatures, and ScaleData, respectively. Dimensionality reduction was achieved via principal component analysis, and the first 20 PCs were used as input for UMAP to visualize cellular relationships. Cell–cell neighborhood graphs were constructed with FindNeighbors, and unsupervised clustering was performed using the Louvain algorithm (FindClusters). Gene expressions of retinal ganglion cells (RGCs) was visualized using FeaturePlot, VlnPlot, and DoHeatmap functions on scaled Seurat data. Marker-positive cell subsets (e.g., *pou4f1⁺*, *pou4f2⁺*, *sncg⁺*, and *isl1⁺* cells) were identified using WhichCells or by subsetting normalized expression matrices. Heatmaps were then generated to display expression patterns within these subsets. Co-expression analysis of *pou4f1*, *pou4f2*, *sncg*, and *isl1* was performed by identifying cells expressing all five genes and quantifying cells expressing individual or combinatorial patterns. The results were visualized using Venn plots and ggplot2 barplots. A similar analysis was conducted for *sncg⁺*, *nefl⁺*, and *nefm⁺* populations, identifying cells co-expressing all three genes and summarizing expression overlaps through the same visualization methods. For comparisons between data sets, after data processing, UMAPs (gene expression) were generated using categorical embedding and continuous embedding functions. Heatmap and violin plot functions in the software were used to create these visualizations. Additionally, trajectory analysis for CiRGCs was performed using the built-in Monocle3-based function in the software. We calculated the root nodes using this function, selected the root nodes within our target cell cluster (anticipated progenitor population), and then ran the function. For comparing multiple scRNA-Seq datasets, we uploaded Cell Ranger-derived raw feature-barcode matrix files for all samples and processed the data simultaneously using the automatic processing functions.

### Single cell ATAC Sequencing

Single-cell ATAC-seq libraries were processed with Cell Ranger ATAC (v2.1.0) to generate fragment, filtered peak-barcode matrices, and metadata. Downstream analysis was performed in R (v4.5.1) using Signac (v1.15.0), Seurat (v5.3.0), SeuratObject (v5.2.0), ggplot2 (*v*3.5.2), dplyr (*v*1.1.4), and GenomeInfoDb (v1.2.14). Filtered matrices were imported with Read10X_h5, integrated with metadata, and only high-confidence nuclei were retained. A chromatin assay was created with CreateChromatinAssay (hg38), and a Seurat object was constructed combining counts, fragments, and metadata. Data were normalized with TF–IDF, and dimensionality reduced via SVD to generate LSI components. UMAP embeddings were computed, neighborhoods identified with FindNeighbors, and clustering performed using the Louvain algorithm (FindClusters. Gene annotations from GRCh38 (Ensembl 109) were imported using rtracklayer (v1.68.0) and GenomicRanges (v1.60.0) and then assigned to the Seurat object. Gene activity scores were calculated with GeneActivity, added as a new assay, and normalized with NormalizeData. Cluster-specific accessibility for RGC marker genes was visualized using FeaturePlot, VlnPlot and DoHeatmap. Cells positive for *POU4F1*, *POU4F2*, *SNCG*, *NEFL*, *NEFM*, or *ISL1* were identified using Which Cells, subsetted, and heatmaps of selected marker genes plotted with DoHeatmap using gradient color scales. Cloupe output from 10X pipeline was visualized using POU4F1+ and POU4F1+RBPMS+ promoter sum positive cells were selected using Loupe Browser 6.5.0 ([Loupe Browser ATAC Tutorial -Software -Single Cell ATAC -Official 10x Genomics Support](https://support.10xgenomics.com/single-cell-atac/software/visualization/latest/tutorial)).a filter function in the algorithm. After selecting these cells, peaks were identified using a peak viewer function. Motif enrichment per cluster was performed in the globally distinguishing mode of the algorithm.

### Pattern ERG and pattern VEP measurement

All animal studies and animal care were performed in accordance with relevant guidelines and regulations approved by IACUC at Children’s Hospital Los Angeles. Both male and female mice were used in all experiments with equal frequency. All the *in vivo* experiments were conducted on 6-8 weeks old mice. For pERG and pVEP recordings, anesthetized animals were placed on a feedback-controlled heated stage set to 37°C. The recordings were conducted using a specific protocol with the default setup from Diagnosys LLC (Boston, MA). 1% tropicamide eye drops were applied to dilate the pupils, and GenTeal tear eye drops were used to prevent corneal desiccation. 6-8 weeks C57/BL6 mice (Jackson lab) were used for baseline measurement. Baseline measurements were taken from 6-8 week old C57/BL6 mice (Jackson Lab). Animals with stable baselines were subjected to NMDA injection (100 mM, 1.5 μl/eye). 7 days after NMDA injection pERG/pVEP was measured to record diminished RGC function (**Fig. 5a**). pERG/pVEP diminished animals were injected with MACS purified CiRGCs/PBS. Following is the brief demonstration of pERG/pVEP measurement: under red light pERG responses were recorded by touching pattern stimulator to cornea of one eye and a full filed stimulator as a fellow eye reference electrode according to manufacturer’s instructions (https://www.diagnosysllc.com/wp-content/uploads/2021/03/Celeris-Pattern-Stimulator-Application-Note-d.pdf). After pERG recordings, the same mice were used for pVEP measurements by adjusting the electrodes to the appropriate channels as per the manufacturer’s guidelines. For pVEP, responses were recorded by placing the pattern stimulator on the cornea, inserting a needle electrode subdermally in the mice’s snout, and positioning the reference electrode at the back of the head. The pVEP protocol was then run using default parameters. Independent pERG and pVEP responses were obtained using the default averaging system. Data, including P1 and N1 amplitude values, were exported from the machine using Espion V6 software (Diagnosys, LLC). Webforms were exported as graphs. Note, NMDA+PBS control group simultaneously used for in another study^36^. We have performed these experiments in parallel.

### Light aversion behavior test

The light/dark apparatus consists of black opaque (100%) acrylic test chambers (30.48 × 15.24 × 30.48 cm (length, width, height)). This chamber was further divided into equal-sized compartments (15.24 × 15.24 × 30.48 cm) by the addition of an insert, to create a dividing wall in the center. Further, to create light and dark zones, one compartment was illuminated with light (2800 lux, similar to cloudy outdoor environment) and the other compartment was kept dark (∼ 0.1 lx). The light and dark compartments were connected by an opening (5 × 5 cm). The position of the mouse within the apparatus was recorded using a camera. Mice were maintained in the testing room for 2 hrs in ambient light (200 Lux) conditions in their home cage with free access to food and water. Each mouse was allowed to habituate to the testing apparatus (both sides) for 10 min while in ambient light. After habituation, one side of the apparatus was illuminated with light at around 2800 lux, and the mouse was allowed to roam freely between each compartment for 5 min. Two parameters, namely the latency of escape from the illuminated area to darkness and the percentage of time spent in the dark using the light intensity, were computed as an index of light sensitivity.

### Visual cliff test

The apparatus was purchased from Conduct Science, Boston, MA, comprises a transparent acrylic chamber measuring 62 cm x 62 cm x 19 cm, featuring a checkered pattern insert. This chamber is divided by a central platform, 1.5 inches high and 2 inches wide, creating two sections. One side is shallow, with a checkered pattern immediately beneath the platform, while the other side is deep, with the same checkered pattern positioned approx. 2 feet below, producing a visual depth illusion. Both sides of the setup are evenly illuminated with a light intensity of approximately 2000-2500 lux. For shallow or deep choice experiments, mice are placed on the central platform, and their movements were captured using a video camera. Each mouse undergoes five trials. In the cumulative duration study, the central platform was removed, and mice were positioned on the midline between both sides. The time spent on each side was then calculated from video recordings. The chamber and platform are thoroughly cleaned after each test. Note, WT and NMDA+PBS control group simultaneously used in another study for both the behavior tests^36^. We have performed these experiments in parallel.

### Statistical analysis and fluorescence quantification

The data is expressed as mean ± standard error of the mean (SEM). Statistical significance was assessed using Student’s t-test, one-way ANOVA, or two-way ANOVA, conducted with GraphPad Prism Software (version 9). The corresponding P value summaries are shown in the figures. For fluorescence quantification, images were analyzed using ImageJ.

## Data availability statement

All data that supports the findings of this study are available from the corresponding author upon reasonable request. Additionally, all sequencing data described in this work has been deposited in the Gene Expression Omnibus (GEO) under accession numbers GSE253807 (SC-RNASeq), GSE253808 (SC-ATACSeq). Upon publication, the GEO will be available for the readers.

---

**Figure S1:**
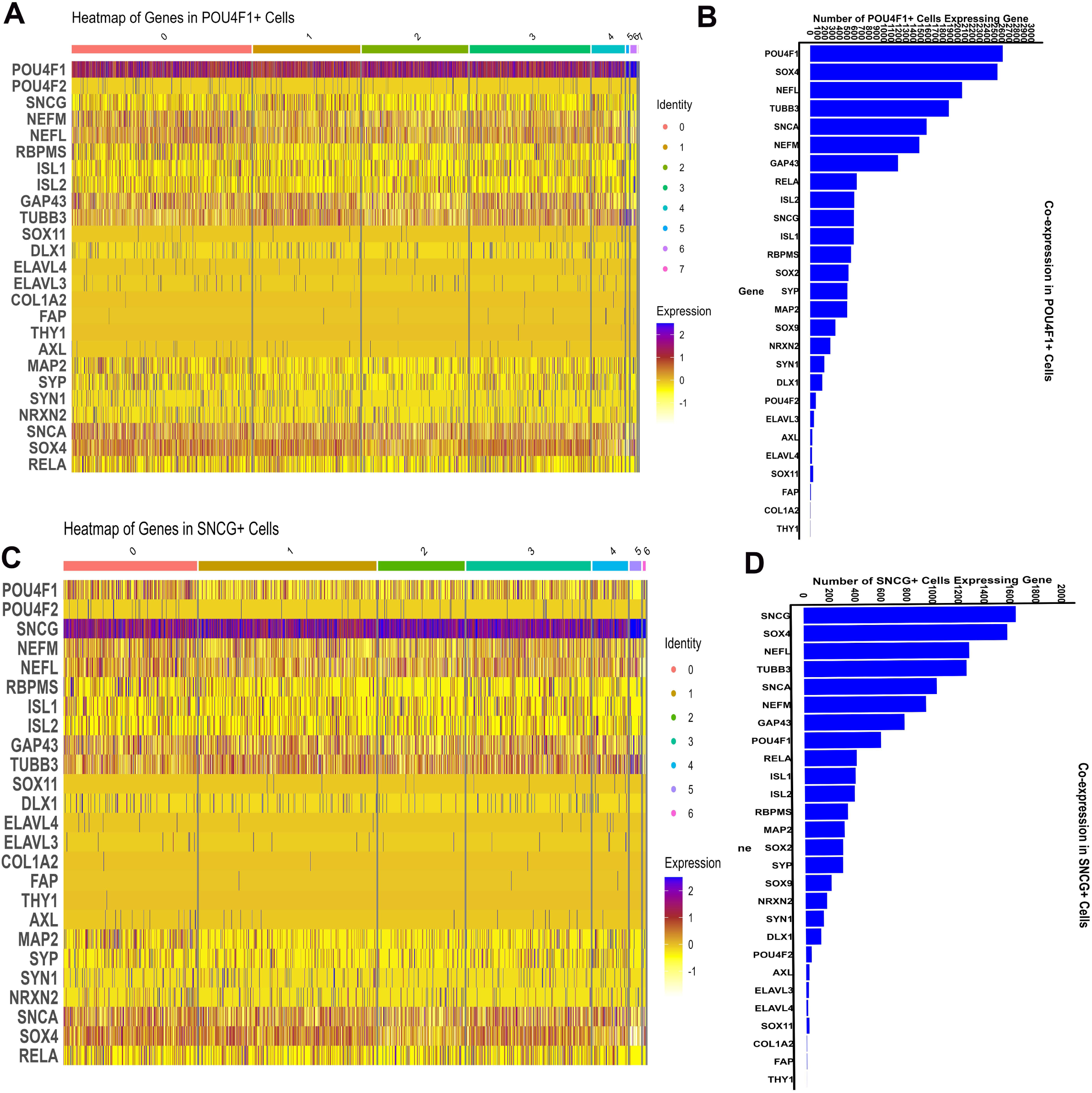
Cell set analysis of open chromatin regions in canonical RGC gene promoters within *Pou4f1*⁺ and *Sncg*⁺ cells. **(A, C)** Heatmaps showing *Pou4f1*⁺ and *Sncg*⁺ cell sets respectively. **(B, D)** Number of cells expressing the indicated genes within the *Pou4f1*⁺ and *Sncg*⁺ cell sets.

**Figure S2:**
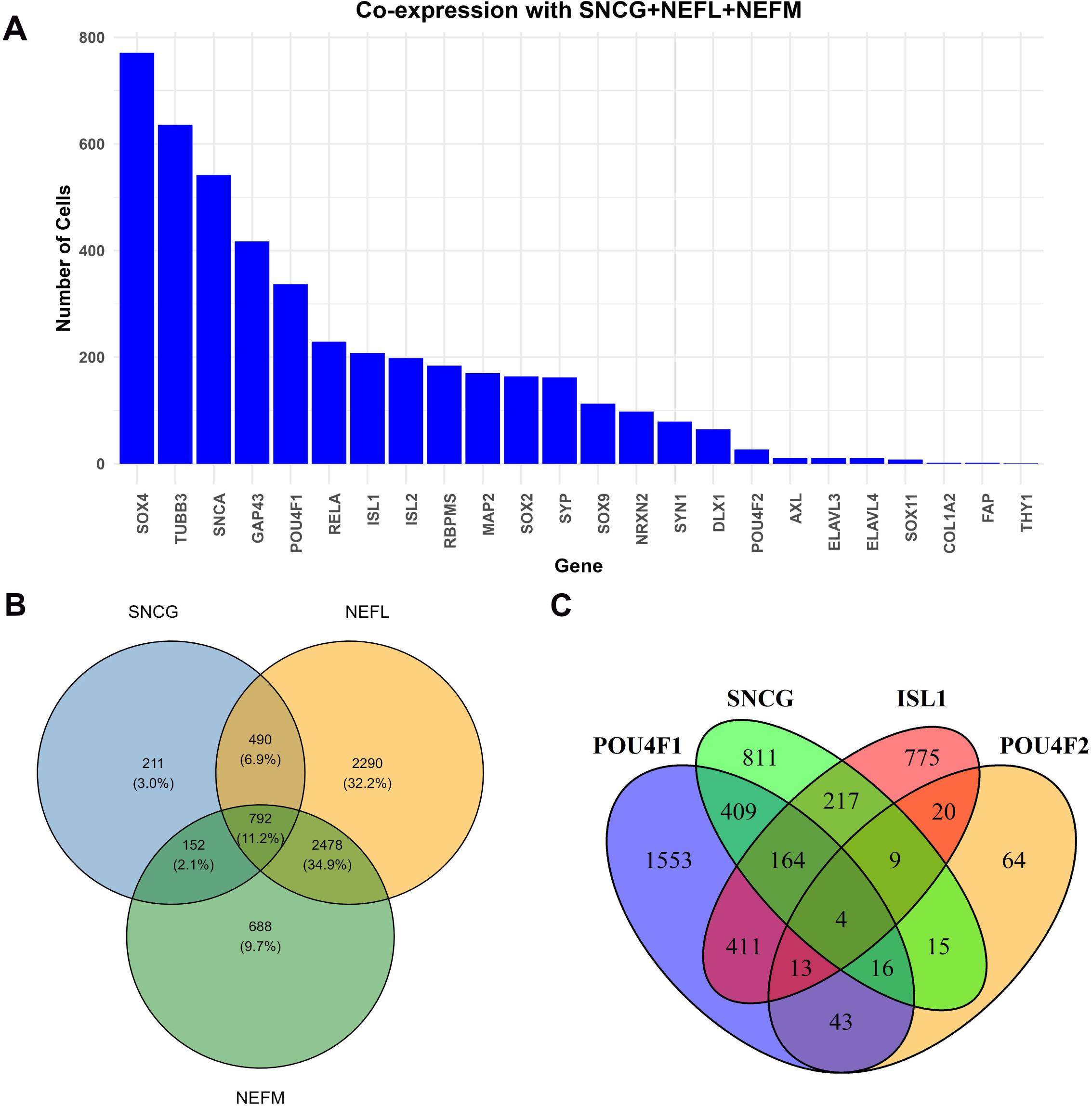
Cell set analysis of open chromatin regions in canonical RGC gene promoters within *Sncg⁺Nefl^+^Nefl^+^* cells. **(A)** Number of cells expressing the indicated genes within the cell sets. (**B**) Pie chart showing the percentage of cells expressing *Sncg, Nefl,* and *Nefm* individually, as well as their co-expression. (**C**) Pie chart showing number of cells expressing *Sncg*, *Pou4f1*, *Pou4f2*, and *Isl1* individually, as well as their co-expression.

**Figure S3:**
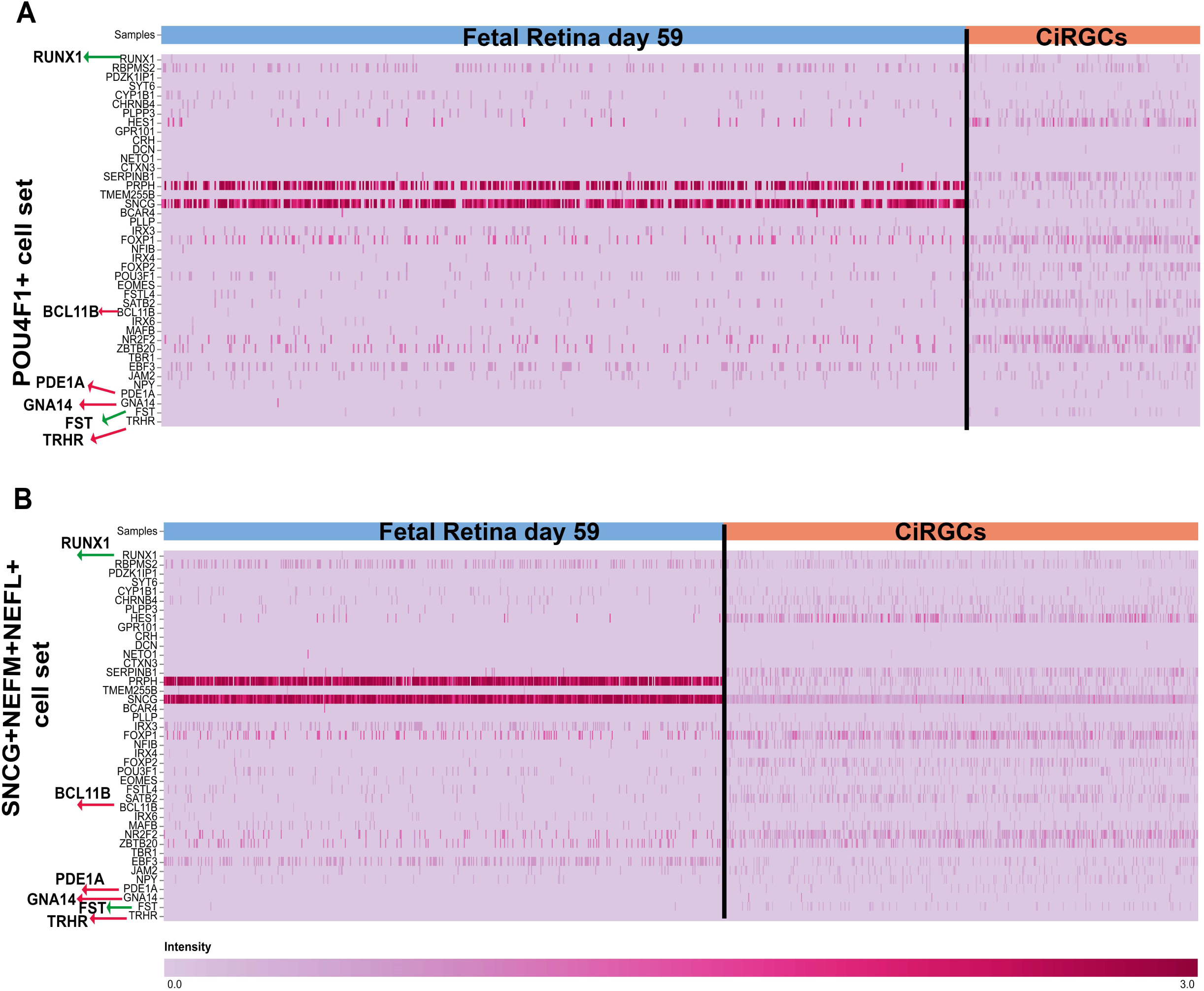
Expression of various RGC subtype specific genes in CiRGCs. Indicated subtypes and corresponding genes are identified based on previous reports.^16^ (**A, B**) Comparisons between CiRGCs (POU4F1+ and SNCG^+^NEFL^+^NEFM^+^ cells) and fetal retina day 59. Red and green arrows are genes showing differential expressions.

**Figure S4:**
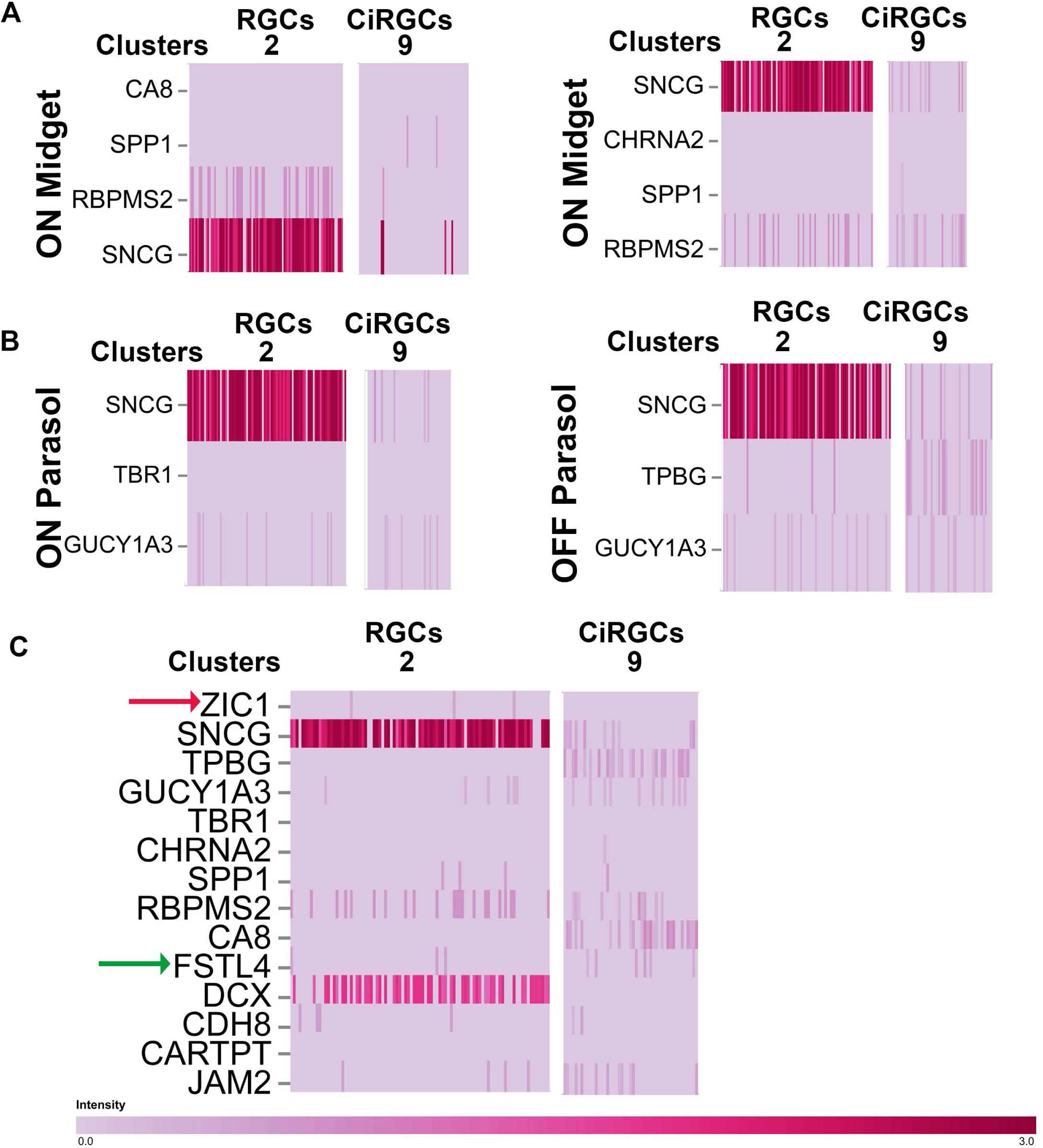
Expression of various RGC subtype specific genes in CiRGCs (POU4F1^+^RBPMS^+^ cell set). **(A)** Cluster wise expression of ON/OFF midget or ON/OFF parasol RGC markers in native RGCs (cluster 2) and CiRGCs (cluster 9) identified in Fig. 2c**. (B)** Expression of indicated subtype genes between same calusters in POU4F1^+^RBPMS^+^ cell set.

**Figure S5:**
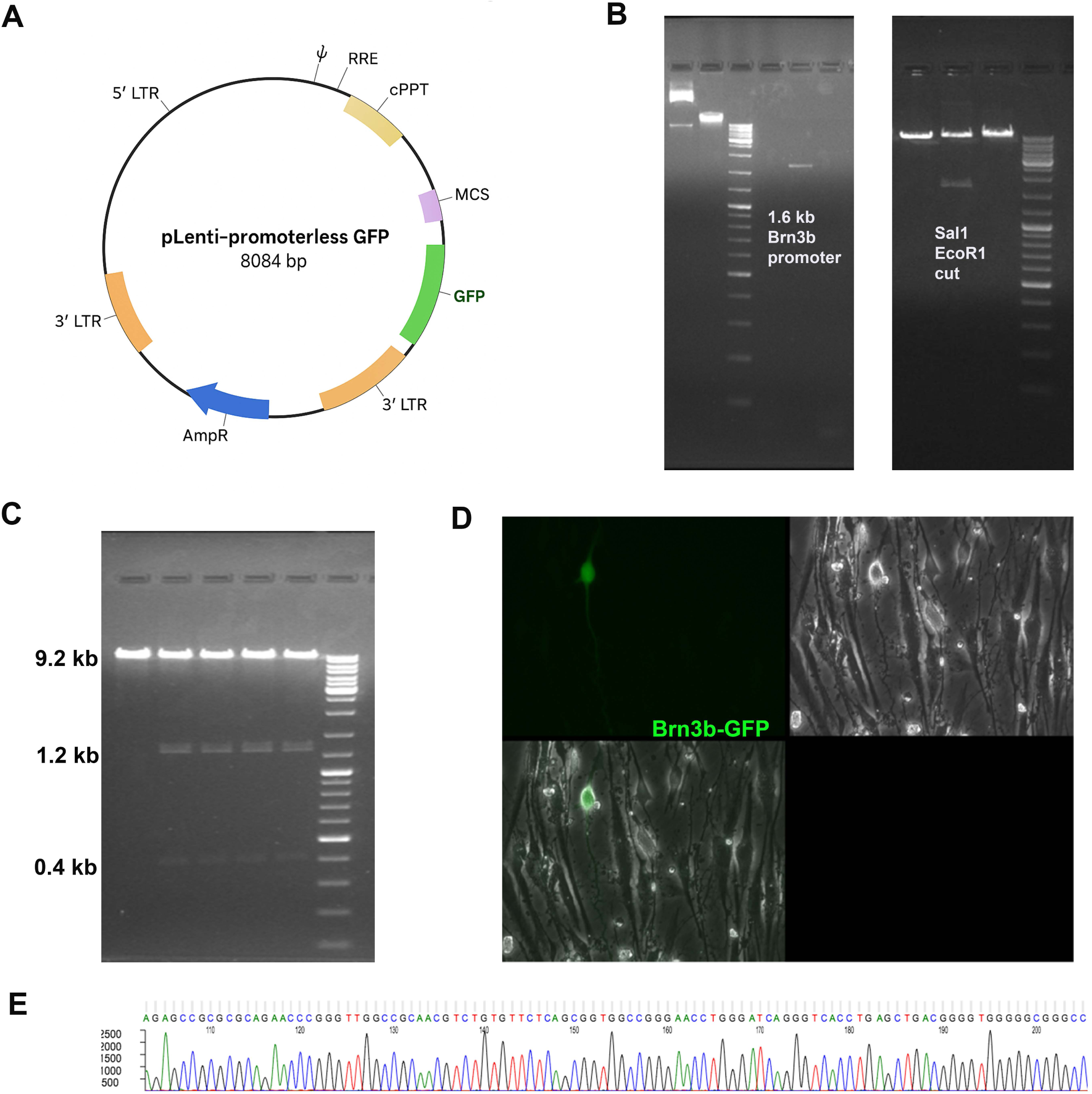
Cloning and sequencing of Brn3b-promoter reporter. **(A)** Map of commercially available pLenti-promoterless-GFP (generated using AI). (**B)** Left: PCR amplification of 1.6 kb promoter segment of Brn3b. Right: DNA fragments showing cloned promoter segment after restriction digestion. **(C)** Restriction mapping of cloned vector using ECOR1 and BamH1 restriction enzymes. (**D**) Brn3b-GFP expression after 5C mediated human fibroblast reprogramming od day 4. **(E)** DNA sequencing of *Brn3b*-promoter segment inside promoterless vector.

**Figure S6:**
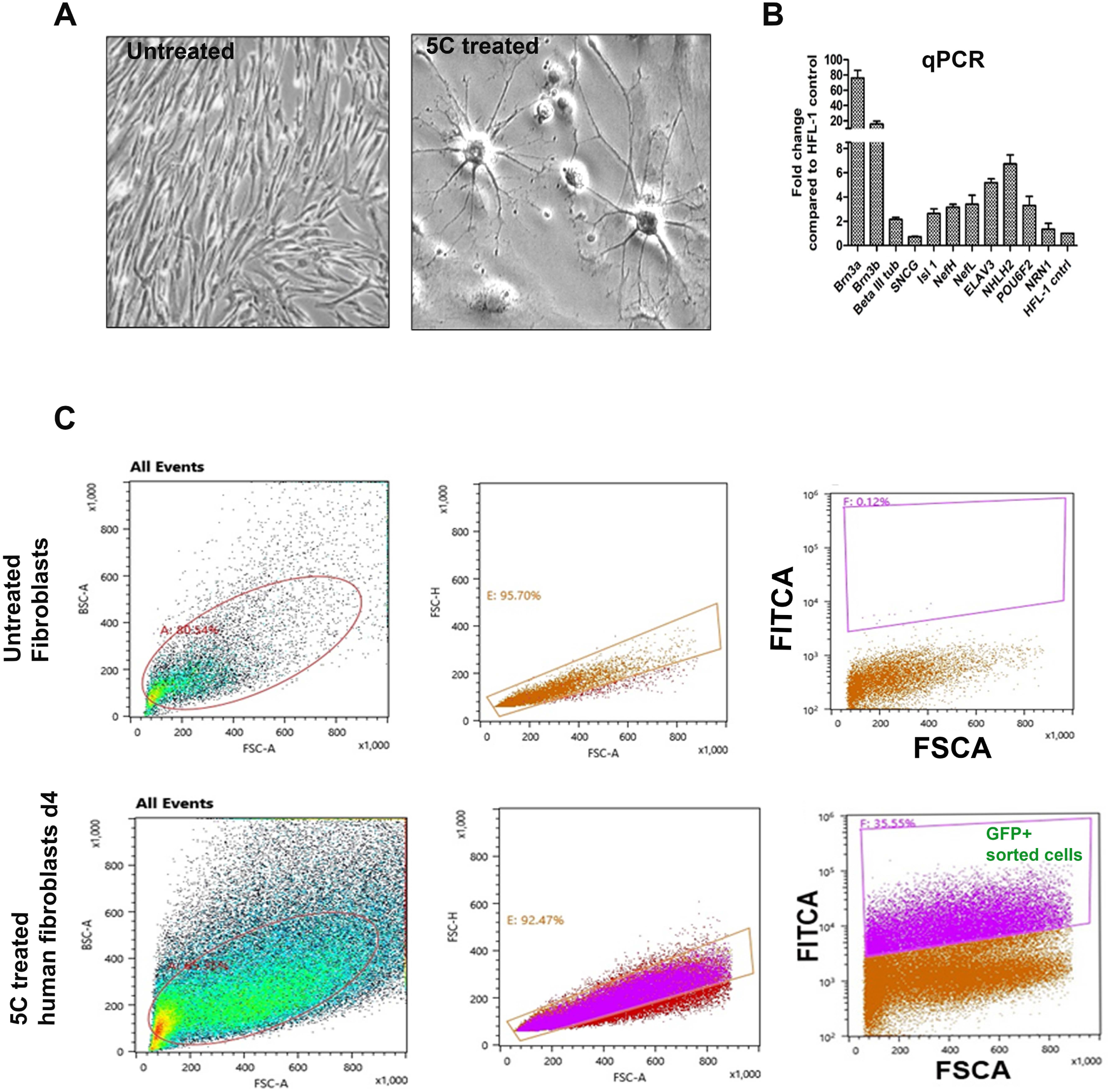
Expression of BRN3b-GFP reporter in 5C mediated reprogrammed CiRGCs. **(A)** Phase contrast image for human fibroblasts and chemically induced CiRGCs. **(B)** qPCR of indicated RGC genes after FACS sorting. **(C)** FACS sorting dot plots showing GFP+ cell separation.

**Figure S7:**
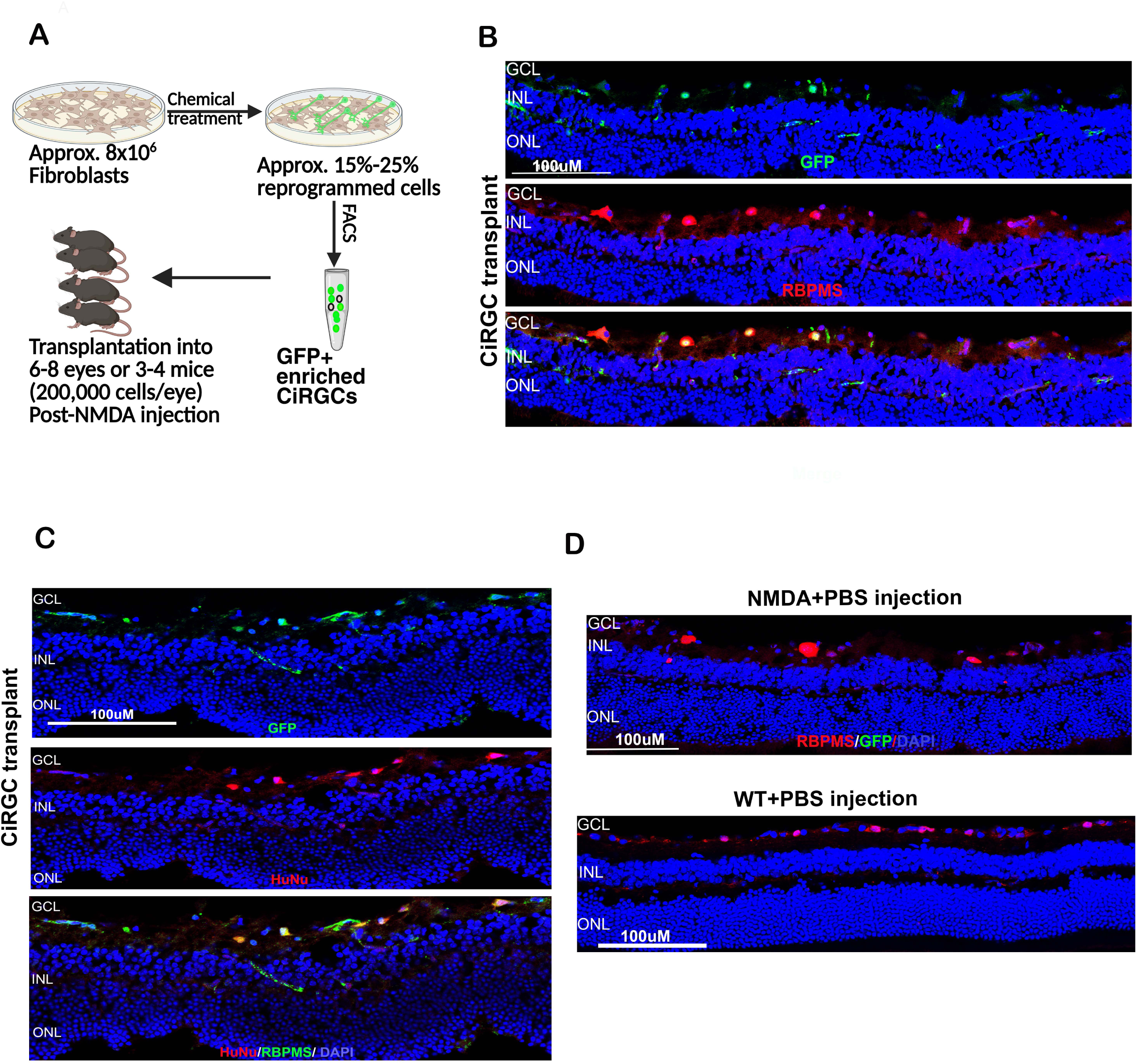
BRN3b-GFP^+^ CiRGCs survival in NMDA injected mice on day 75. **(A)** Scheme for the transplantation experiment. **(B)** Images showing CiRGC and PBS injected retina stained with RBPMS and anti GFP antibodies. **(C)** Human nuclear antigen (HuNu) staining of transplanted CiRGCs. **(D)** Survived CiRGCs in a separate co-hort 4 weeks after transplantation. a-c; MACS purified cells and d, FACS purified cells were used for transplantation.

**Figure S8:**
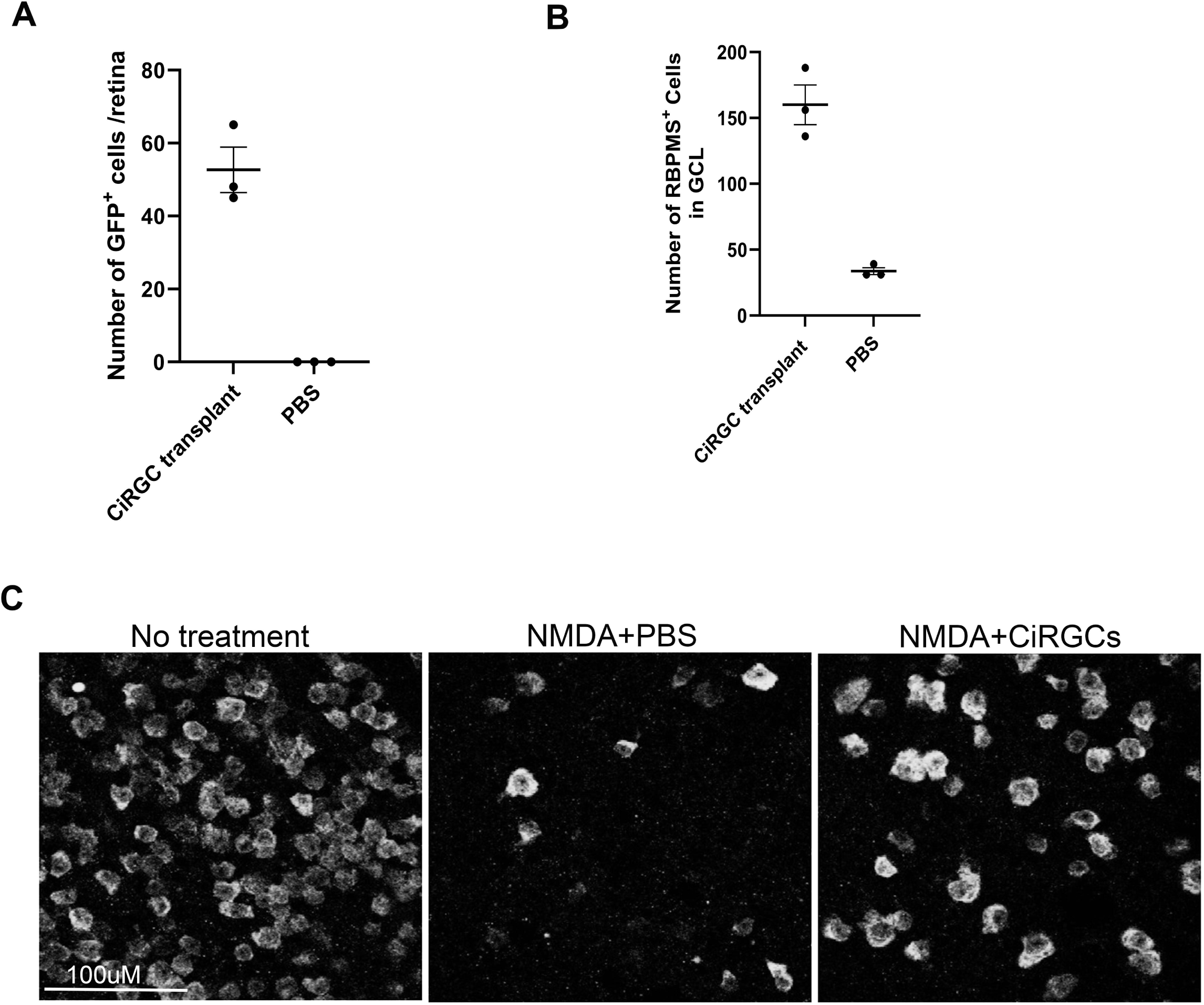
Quantification of CiRGC survival and neuroprotection. **(A)** Quantification for number of GFP+ CiRGCs per eye on day 75**. (B)** Quantification of number of survived RBPMS+ RGCs per eye on day 75. **(C)** representative images for RBPMS cell count indicated in B. For “A” n=3 eyes, 4 sections/eye, 3 fields/section were counted. For “B” n=3 eyes were counted.

**Suppl. Table 1:** Retinal Differentiation Medium 2 (RDM2) Composition.

| **CiRGC Differentiation Medium** |  |  |  |
| --- | --- | --- | --- |
| **Components** | **Stock Conc.** | **Final Conc.** | **Volume added in 50 mL** |
| Knockout Serum Replacement | 100 % (regular stock) | 10% | 5 ml |
| B27 Supplement | 100x | 1% | 500 µl |
| N2 Supplement | 100x | 1% | 500 µl |
| Noggin | 20 µg/ml | 10 ng/ml | 25 µl |
| DKK1 | 10 µg/ml | 10 ng/ml | 50 µl |
| IGF-1 | 200 µg/ml | 50 ng/ml | 12.5 µl |
| bFGF | 100 µg/ml | 5 ng/ml | 2.5 µl |
| SB431542 | 50 mM | 5 µM | 5 µl |
| DMEM/F12 with Glutamax |  |  | Up to 50 ml |
| **RDM1 medium composition** |  |  |  |
| RDM2 minus SB431542 |  |  |  |

**Suppl. Table 2:** Antibodies used for this study.

| **Name of Antibody** | **Host** | **Dilution** | **Vendor** | **Cat#** |
| --- | --- | --- | --- | --- |
| Anti-Brn3a | Mouse | 1:100 | Santa Cruz | SC-8429 |
| Anti-RBPMS | Rabbit | 1:500 | GeneTex | GTX118619 |
| Anti-SNCG | Mouse | 1:500 | Abnova | H00006623-M01 |

**Suppl. Table 3.**
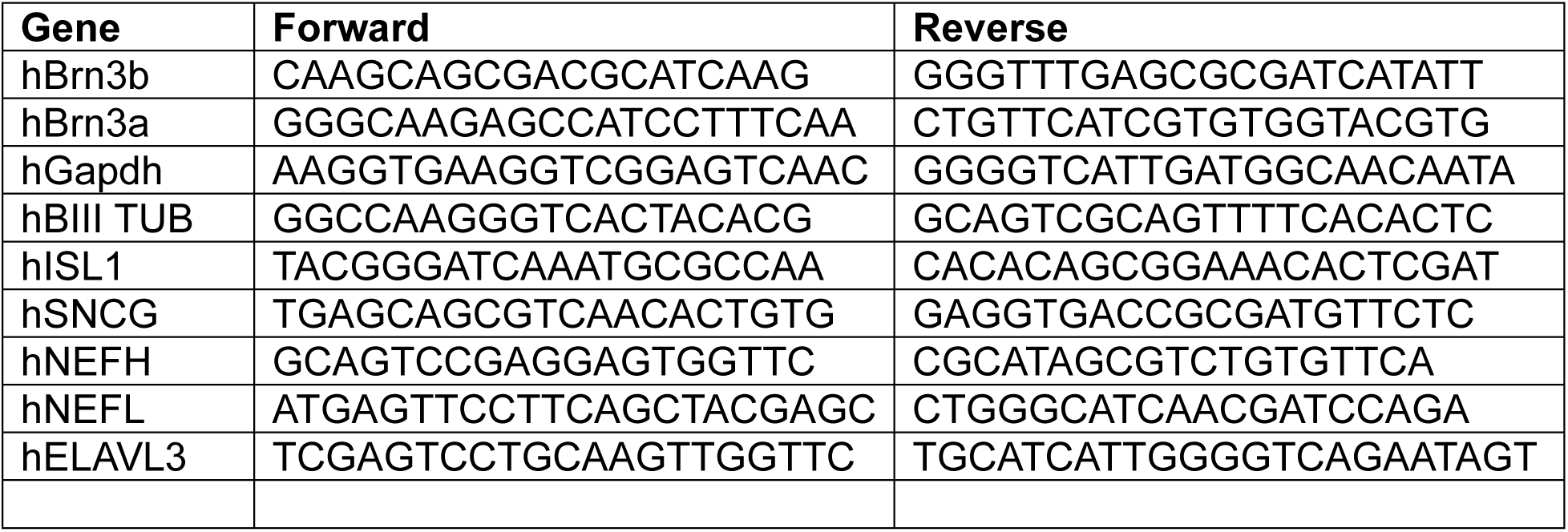
Primer lists.

## Notes

### Competing Interest Statement

The authors have declared no competing interest.

### Funding

- Glaucoma Research Foundation
- Knights Templar Eye Foundation
- California Institute for Regenerative Medicine
- Children’s Hospital of Los Angeles

